# Erroneous compensation for long-latency feedback delays as origin of Essential Tremor Running title: Erroneous delay compensation as origin of ET

**DOI:** 10.1101/2024.01.11.575204

**Authors:** Flo Blondiaux Pirson, Lise Colmant, Louisien Lebrun, Bernard Hanseeuw, Frédéric Crevecoeur

## Abstract

Essential tremor (ET), a movement disorder characterized by involuntary oscillations of the limbs during movement, remains to date not well understood. It has been recently suggested that the tremor originates from impaired delay compensation, affecting movement representation and online control. Here we tested this hypothesis directly with ET patients (N=24) and neurologically intact (NI) volunteers (N=28) in an upper limb postural perturbation task. After maintaining their hand in a visual target, participants experienced perturbations of unpredictable direction and magnitude, and were instructed to counter the perturbation and steer their hand back to the starting position. In comparison with NI volunteers, ET patients’ early muscular responses (Short and Long-Latency Responses, 20-50 ms and 50-100 ms respectively) were preserved or even slightly increased. However, they exhibited perturbation-dependent deficits when stopping and stabilizing their hand in the final target supporting the hypothesis that the tremor was generated by the feedback controller. We show in a computational model that errors in delay compensation accumulating over time produced the same small increase in initial feedback response followed by oscillations that scaled with the perturbation magnitude as observed in ET population. Our experimental results therefore validate the computational hypothesis that inaccurate delay compensation in long-latency pathways could be the origin of the tremor.

**Significance Statement:** Essential Tremor origin remains poorly understood. In the present study, we focused on motor impairments associated with feedback control. Following a mechanical perturbation applied to their arm, patients’ short and initial long-latency stretch responses were preserved. However, we observed clear impairments during the stabilization phase that scaled with the perturbation magnitude. These results were reproduced in a computational model where delay compensation was inaccurate, suggesting that the origin of the tremor may lie in an underestimation of the delays impacting the internal monitoring of the motor commands.

**Conflict of interest:** The authors declare no competing financial interests.

## Introduction

Essential Tremor (ET), among the most frequent movement disorders (Louis & Ferreira, 2010), typically manifests as involuntary oscillations of the upper limbs and less frequently head, legs and trunk, mainly during movement (Bhatia et al., 2018; Shanker, 2019). While tremor characteristics, such as frequency and lack of impact of loading, have been described extensively, the underlying pathophysiology remains poorly understood (Elble, 2003). Several clinical, postmortem and neuroimaging studies pointed out that ET could originate from the cerebello-thalamo-cortical loop (Ibrahim et al., 2021), without characterizing how the movement deficits arise from a functional perspective. Identifying preserved and impaired cerebellar functions in ET would therefore help to better understand the mechanisms at the origin of the tremor (Butler et al., 2023; Louis & Faust, 2020b; Trillenberg et al., 2006).

From a computational perspective, it is well known that the presence of delays in a feedback control system can induce oscillations if the impact of the delay is not taken into account (Miall et al., 1993; Stein & Oĝuztöreli, 1976). This parallels with the way we control our limbs. Indeed, the sensory feedback, used in motor control, is delayed due to the conduction time of sensory signals and motor commands through the nerves. The delay depends on the feedback loop: in the upper-limb, short-latency delays associated with spinal stretch reflex are in the range of 20-30 ms, and long-latency delays involving a transcortical loop are in the range of 50-60 ms (Scott, 2016). Importantly, it has been also suggested that state estimation, and therefore compensation for temporal delays, is performed in the long-latency feedback pathway (Crevecoeur & Scott, 2013). Assuming erroneous delay compensation in this sensorimotor system, previous simulations reproduced oscillations in the frequency range observed in ET (Crevecoeur & Gervers, 2019), allowing us to formulate the hypothesis that ET originates from errors in the long-latency feedback loop.

Indeed, Long-Latency Responses (LLRs) refer to the burst of muscular activity observed 50-100 ms (for upper-limbs) post-perturbation (Pruszynski & Scott, 2012). This activity helps to counteract perturbations and is likely supported by transcortical feedback. LLRs are not only elicited for large perturbations but are also present for very small perturbations, with a magnitude close to the magnitude of errors observed during voluntary control (Crevecoeur et al., 2012), and parallels unperturbed movement in the tendency to exploit target redundancy (De Comite et al., 2021). This suggests that LLRs could therefore be present constantly during movements to compensate for the undesirable effects of motor variability. If LLRs continuously contribute to movement control, errors in this pathway could lead to general movement deficits affecting both voluntary movements and feedback responses to perturbations. This hypothesis is also supported by our previous study showing that ET patients could learn to anticipate the effect of a perturbation but exhibited adaptation impairments linked to online control (Blondiaux et al., 2023). Together, there is strong evidence that movement execution is impaired in ET, and errors in the long-latency feedback loop is a candidate model for their deficits.

We tested this hypothesis directly by investigating feedback control in both ET patients and a neurologically intact (NI) control group using a postural perturbation task. We observed similar initial perturbation-related limb displacement and intact short (R1: 25-45 ms post-perturbation) and initial LLR (R2: 45-75 ms), suggesting preserved peripheral system and initial stretch responses. However, perturbation-dependent deficits arose during the later phase of the feedback correction and during stabilization, which followed an enhanced stretch response in the late LLR epoch (R3: 75-105 ms) dependent on the load magnitude. We reproduced both movement kinematics and transient perturbation-related increase in the stretch response in a computational model considering an error in the delay parameter. Together our analyses suggest that ET could arise from partial or inaccurate compensation of long-latency delays, leading to oscillations in movements caused by accumulation of error over time.

## Methods

### Participants

A total of 24 ET patients (14 females, mean age: 61.1 ± 13.1) and 28 NI volunteers (17 females, mean age: 62.7 ± 12.6) were recruited for this study. The ethics committee at the host institution (*Comité d’Éthique Hospitalo Facultaire*, UCLouvain, Belgium) approved the experimental protocol and all participants provided written informed consent following standard procedures. All participants were evaluated using the Fahn-Tolosa-Marin Tremor Rating Scale (FTM-TRS) and participants with other neurological disorders were excluded. The FTM-TRS scale was divided into 3 sections. Part A quantified the tremor amplitude of the limbs at rest, during postural control, and during action. Part B assessed writing, drawing and pouring liquids. Finally, Part C evaluated the impact of the tremor in daily life activities (eating, drinking, getting dressed, working, …) (Fahn et al., 1993). The latter part was filled by the participant and was not considered during this study due to its inherent subjectivity. ET patients did not interrupt their medication prior to the evaluation. The FTM TRS Score for part A and B was 2.3 ± 0.38 (mean ± SEM) for the NI group and 21.6 ± 1.98 for the ET group.

### Task

Participants sat on a height-adjustable chair and grabbed the robotic handle of an End-Point KINARM device (KINARM, Kingston, Ontario). Direct vision of volunteers’ hands was blocked and a semitransparent mirror that reflected the virtual reality screen (VPixx, 120 Hz), allowing them to interact with virtual targets (Figure 1a). A hand-aligned cursor was always represented (white dot, radius 0.6 cm). The experiment was composed of 2 trial types: voluntary movements and perturbation trials.

**Figure 1:**
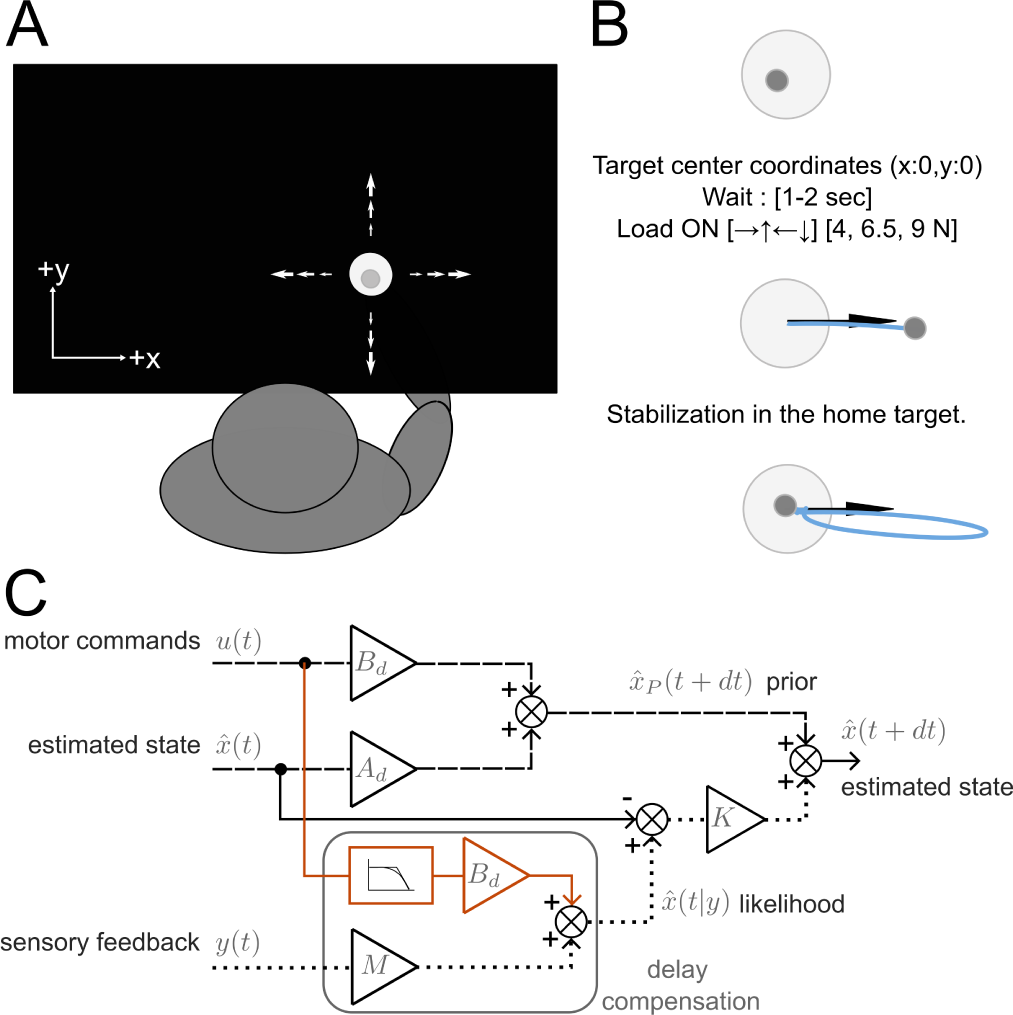
S*c*hematic *representation of the protocol design and theoretical model*. **A**, Prior perturbation, participants maintained the hand-aligned cursor in the home target. Direct view of the limb was blocked by an occluder but participants had continuous visual feedback about the hand-aligned cursor. **B**, Perturbation trials started in the home target, participants received one of three perturbation magnitudes (4, 6.5, 9N) in one of four directions (+x,-x,+y,-y). Visual feedback about movement timing was provided via target color (see methods). **C**, Model pathway adapted from Crevecoeur and Gevers (2019). Model compensating for the delay in sensorimotor control. The dotted pathway corresponds to the sensory feedback, the dashed pathway corresponds to the internal prior and the orange pathway correspond to the delay compensation that is suspected to be faulty in ET. 𝐴_𝑑_ and 𝐵_𝑑_ define the dynamics of the model, dt is the time step, 𝛿𝑡 the system delay and K are the Kalman gains. For clarity, we omitted the explicit dependencies of M on 𝛿𝑡 and K on t, see methods for variable and matrices definitions.

Participants started with a block of 60 voluntary movements. A home target of radius 1 cm was presented approximately 30 cm ahead of the participant’s torso, aligned with the right hand. They had to maintain the hand in the home target (coordinates: [x, y] = [0, 0]), after a random time uniformly distributed between 1 and 2 sec, an external target was lit and used as a go signal. Participants were instructed to pass through the external target (radius 2.5 cm) and to finish their movements inside the home target. The goal target was randomly selected within 4 targets with coordinates [10,0], [-10,0], [0,7.5], and [0,-7.5]. Each target was presented 15 times during the block. Participants were encouraged to finish their movements in less than 2 s. This initial block with voluntary movements was performed to collect information about movement kinematics without perturbations, but the results were consistent with the main experiment and with previous reports on reaching movements in this population (Blondiaux et al., 2023). Because these results did not provide specific additional information, we do not show the corresponding dataset here.

Perturbation trials started with the same home target (Figure 1B): participants maintained the hand in the home target, after a random time between 1 and 2 s, the robot applied a step force (rise time: 5 ms) in one of 4 directions [+x,-x,+y,-y], with a magnitude of 4, 6.5, or 9 N. The perturbation magnitude and direction were randomly selected. The participants performed a total of 300 trials composed of 25 trials per combination of perturbation direction and magnitude. The trials were divided into 5 blocks of 60. Participants were instructed to return and stabilize in the home target, and they received a positive mark indicating good performance (target turned green) if they completed their movement in less than 3 s. The total score was projected on the screen for their motivation. When movements were too slow, the target remained red. Trial timed out after 3 s if the participant was not able to stop their hand. The criterion on movement time was applied to encourage consistent speeds, but all movements that returned to the target were kept for analyses. The step force turned off at the end of the trial and the inter-trial time was 1 s. Participants were encouraged to take short breaks between the blocks to reduce fatigue (15 s to 2 min).

For the perturbations along the y-axis, we did not observe any significant scaling of the EMGs with perturbation magnitude both in the NI and ET group. This was likely due to participants’ limb configuration (the arm and forearm were approximately vertical and horizontal, respectively) and of the action of gravity, which in this direction passively brought the hand back to the initial position. Thus, although all perturbations were applied during data collection to reduce anticipation, the following analyses focus on the positive and negative perturbations along the x-axis, for which we collected a clear EMG response that scaled with the perturbation magnitude.

### Data collection and analysis

Kinematics data was filtered using a dual pass 4^th^ order low pass Butterworth filter with a cut- off frequency of 20 Hz. Movement velocity was extracted using KINARM’s data extraction scripts (version 3.1) and movement path lengths were defined as the integral of the velocity vector norm. Movement onset and end were defined using a speed threshold of 3 cm/s for at least 200 ms. Movements were included in the dataset if the participant returned to the target after perturbation. Movements oscillating around the target at the end were also included in the analysis when possible. Based on these criteria, we kept all trials for ET participants and removed 1 trial for two NI volunteers who did not return to the target. NI volunteers managed to stop all their trials at the end of the trial time, while on average, ET participants managed to stop for a fraction of trials corresponding to 89.42% ±4.36% of the total number of trials. The return time was defined as the duration between perturbation onset and the moment when the hand re-entered the target. The stabilization phase was defined as the remaining duration of the trial. The Power Spectral Density (PSD) was computed on the acceleration signal during the stabilization phase, and averaged across perturbation trials of similar direction and force magnitude. Subsequently, the average PSD for each participant was averaged across populations (NI and ET), and normalized to the peak PSD observed in ET patients.

Surface EMG electrodes (Bagnoli surface EMG sensor, Delsys INC. Natick, MA, USA) were used to record the muscular activity during movement. The 6 electrodes were located on the Pectoralis Major (PM), Posterior Deltoid (PD), Biceps, Brachialis, and Long and Lateral heads of the Triceps. The electrodes were attached after light abrasion of the skin with cotton and ether. A conductive gel was used to improve data quality. The EMG data was sampled at a frequency of 1000 Hz. The reference electrode was located on the right ankle. Raw EMGs were filtered using a dual pass 6^th^ order Butterworth filter with a passband frequency range of 20 Hz to 250 Hz, rectified and aligned on the perturbation onset for analyses. To normalize EMG data, participants had to apply a force of 15 ±1 N during 3 s on the handle which was locked by the robot for this calibration procedure. A force gauge was displayed on the screen to help applying the target force. Calibration blocks of 12 trials (3 trials in each direction: ±𝑥, ±𝑦 (unused)), were used at the beginning, middle and end of the experiment (3 blocks in total). We normalized the EMG of each muscle by dividing the signals by the mean value across the calibration trials.

### Statistical Analysis

Comparisons across ET and NI populations were performed based on linear mixed models (LME). The fixed predictors were the perturbation *Direction* (2 level categorical factor, -x or +x) and *Force* (numerical) as well as the *Diagnosis* (2 level categorical factor, NI or ET), plus a random intercept to capture inter-individual variability. We use 𝑌_𝑖𝑗_ to designate the dependent variable of interest, with index *i* corresponding to the trial number and *j* to the participant. We used a significance threshold of 0.05 given the higher variability inherent with clinical population, as well as with the between-participants design. LME were used for the kinematics analyses with 𝑌_𝑖𝑗_ representing respectively for the movement phase the pre-perturbation average speed, displacement at 100 ms, maximum displacement, path length and return time and for the stabilization phase, the stop time, maximum displacement, and path length. LME were also used in the EMGs analyses for the average EMG activity prior the perturbation (-50-0 ms), short latency (R1:25-45 ms), and long-latency bins (R2:45-75 ms and R3:75-105 ms). Effects of the direction and the force (p-value smaller than 10^−3^in all tests) will not be reported. The statistical model can be written as follow:

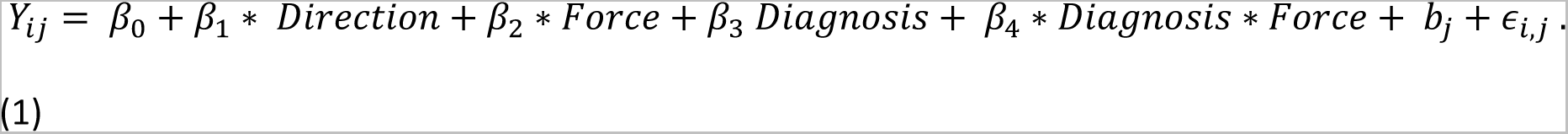

### Model

We reimplemented the model described in Crevecoeur and Gevers, and refer the reader to this reference for a fuller description (Crevecoeur & Gervers, 2019). The simulations describe feedback control of a single joint system (Figure 1C). We considered three torque loads acting on the system: a dissipative torque opposite and proportional to the velocity with a constant G = 0.14 Nm/rad/s (Crevecoeur & Scott, 2014) , a controlled torque T and an external torque 𝑇_𝑒𝑥𝑡_. We considered a first-order filter as linear model of the muscle, transforming the command 𝑢(𝑡) into the control torque with a time constant 𝜏 = 60 𝑚𝑠. The system dynamics is captured by the following system of continuous differential equations:

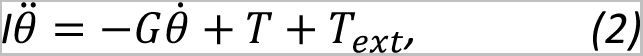

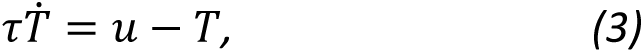

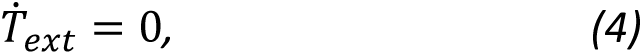

with the inertia set to 𝐼 = 0.15 Kgm², the joint angle 𝜃, and dots represent time derivatives. We defined the state vector 𝑥(𝑡) = [𝜃 ^𝜃̇^ 𝑇 𝑇_𝑒𝑥𝑡_]^𝑇^ and rewrote the three equations as follows:

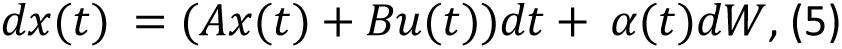

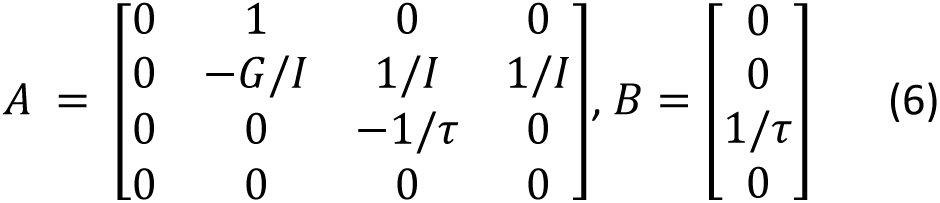

The sensory feedback was defined as follows:

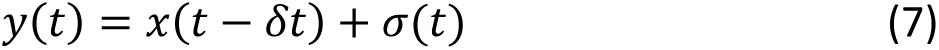

In Equation 5 and 7, 𝛼(𝑡)𝑑𝑊 represented signal-dependent and additive sources of noise, 𝑦(𝑡) was the current sensory feedback signal, 𝑥(𝑡 − 𝛿𝑡) the delayed state with 𝛿𝑡 = 55 𝑚𝑠, and the sensory noise 𝜎(𝑡)∼𝑁(0, Σ_𝜔_). We augmented the state with the target coordinates 𝑥^∗^(𝑡). We then discretized the system using a discretization step of 𝑑𝑡 = 5 𝑚𝑠. The system became:

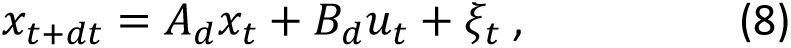

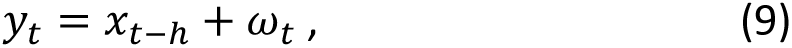

Where 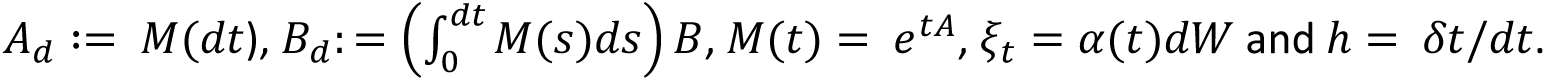

Because of delayed and noisy sensory feedback, we needed to compute an estimate of the state vector:

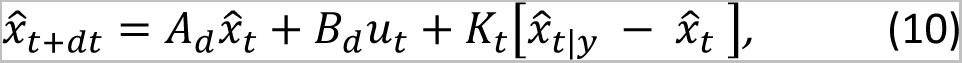

where we define the prior estimate as 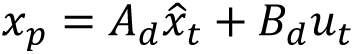. The Kalman gains 𝐾_𝑡_ were computed using a forward recursion.

This formulation differs from the standard definition of predictive Kalman filter, by incorporating an explicit extrapolation of the system state given present sensory feedback. Indeed, the standard form of Kalman filters uses an *innovation sequence* for the non-delayed case equal toy_t_ − 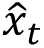. Here, because 𝑦_𝑡_ is a delayed state measurement, it is possible to extrapolate this information and compute the current expected state given 𝑦_𝑡_, the knowledge of limb dynamics and the motor commands during the delay interval. This extrapolation is either performed recursively based on system augmentation, or explicitly in the formulation that we have chosen here which corresponds to finite spectrum assignment (Mondie & Michiels, 2003). With the discrete time system equations, the extrapolation of delayed sensory signals to the present state takes the following form:

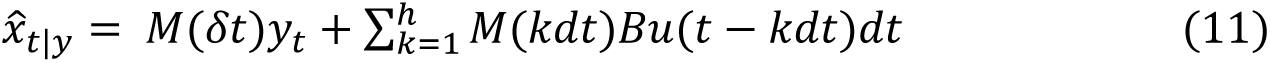

In the sequel, we assume that this operation is performed with a value of the delay 𝛿𝑡 that underestimated the correct value. Recall that, in addition to the explicit dependency of 𝛿𝑡 on M, the motor commands were integrated over a smaller delay interval, h being the delay in discretized time steps. Observe this filter is equivalent to a standard Kalman filter when the delay 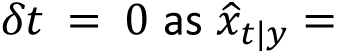. It is also similar to system augmentation (Crevecoeur & Scott, 2013; Fridman, 2014), where the estimation of the system state during the delay interval results from the augmented structure of the state vector. It is important to emphasize that we assumed an output signal of the form of Equation 9. If a more general output signal in the form 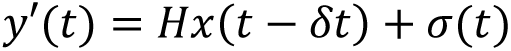 must be considered, then it is possible to reconstruct an estimate of the state vector at 𝑡 − 𝛿𝑡 with an observer prior the extrapolation step. Such intermediate observer would produce an estimate of the state that corresponds to Eqn. 9.

In order to calculate the control signal, we minimized a quadratic cost function penalizing position and velocity error (first term) and control (second term), which writes as follows:

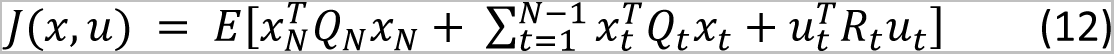

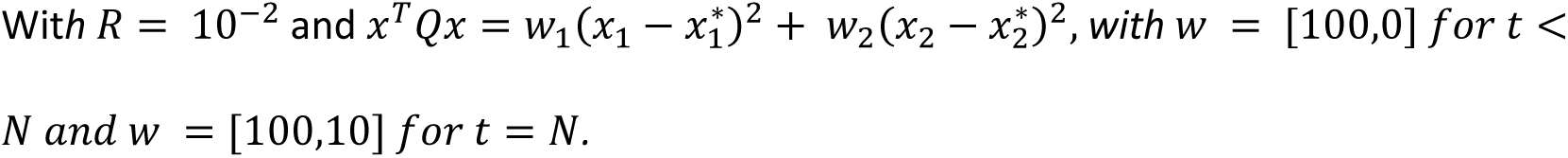

Based on these assumptions, the optimal motor commands were defined as 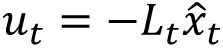 The optimal feedback gains 𝐿_𝑡_ were obtained via a backward recurrence following standard technique.

For the simulations, to capture errors in the delay estimation, we considered a case for which the delay (𝛿𝑡 = 55 𝑚𝑠) was underestimated. We used 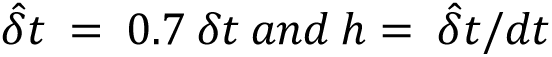 in Equation 11, with ℎ rounded to the closest integer. A delay error of 70% was used to match qualitatively the experimental data and verified in a sensitivity analysis that our conclusion did not depend on this choice. For the simulations with noisy underestimation of the delay we added or removed up to 10ms to the estimated delay. The noise was randomly and uniformly selected from the set {-10, 5, 0, 5, 10}ms. For the simulations with an error in the prior estimate (Eqn. 10), we multiplied it by a factor larger or lesser than 1 (1.1 and 0.7, respectively). Note that this is not equivalent to multiplying 𝐴_𝑑_ and 𝐵_𝑑_ by a scaling factor, as these matrices are also used in the delay compensation step (Figure 1C). The step perturbations amplitudes were 1, 2 and 3 Nm applied at 50 ms. Movement duration was of 800 ms. The figures were produced using the average across 20 simulations in each condition. Similar to the data, the PSD of simulated traces was computed on the acceleration during the stabilization phase and averaged over trial repetitions then perturbation magnitude. Then, the PSD was normalized using the maximum PSD value from the simulations obtained with an error in the delay. An access to working code for the model simulations is provided at the following URL: []. The experimental dataset can be accessed at [].

## Results

We studied the ability of ET patients (n=24) and NI volunteers (n=28) to respond rapidly to step perturbations of various directions and amplitudes. After holding the handle of a robotic arm in the initial visual target, participants were instructed to counter the perturbations and steer their hand back to the initial target (Figure 1). Four directions and three force magnitudes were used for this experiment with a total of 300 randomly interleaved trials (25 repetitions per direction and magnitude). The analyses focus on the lateral perturbations (aligned with positive and negative values along the 𝑥-axis, 150 trials, see methods).

Figure 2 A-C provides an overview of the hand traces of an exemplar participant of each group for each perturbation. Initially, the perturbation had a similar effect on the ET participants, however, once participants returned to the target and tried to stop their movements, oscillations around the target became visible. We observed a slightly larger pre-movement speed explained by oscillations within the target in some heavily affected patients, resulting in a significant effect of the diagnosis (Figure 2D; LME: Diagnosis: F(50) =2.39 , P=0.02). We decomposed movements in two phases: movement and stabilization, with the cut-off being the first time the participant re-entered the target. Concerning the movement phase, the perturbation initially impacted ET patients and NI volunteers in a similar way. There were no significant differences in hand displacement at 100 ms (Figure 2E; LME: Diagnosis: F(50) =0.32, P=0.75; Diagnosis x Force: F(7743) =1.68, P=0.09), in maximum hand displacement (Figure 2F; LME: Diagnosis: F(50) =0.29, P=0.78; Diagnosis x Force: F(7743) =-1.79, P=0.07), in path length (Figure 2G; LME: Diagnosis: F(50) =0.25, P=0.80; Diagnosis x Force: F(7743) =-1.05, P=0.29) and return time (Figure 2H; LME: Diagnosis: F(50) =-0.46, P=0.65; Diagnosis x Force: F(7743) =0.11, P=0.92). However, during the stabilization phase, ET participants faced clear difficulties, with a longer stopping time (Figure 2I; LME: Diagnosis: F(50) =2.83, P<10⁻³; Diagnosis x Force: F(7362) =5.05, P<10⁻⁴), larger oscillations defined as maximum displacement during the stabilization phase (Figure 2J; LME: Diagnosis: F(50) =0.44, P=0.67; Diagnosis x Force: F(7743) =3.66, P<10⁻³) and increased path length during that phase (Figure 2K; LME:

**Figure 2:**
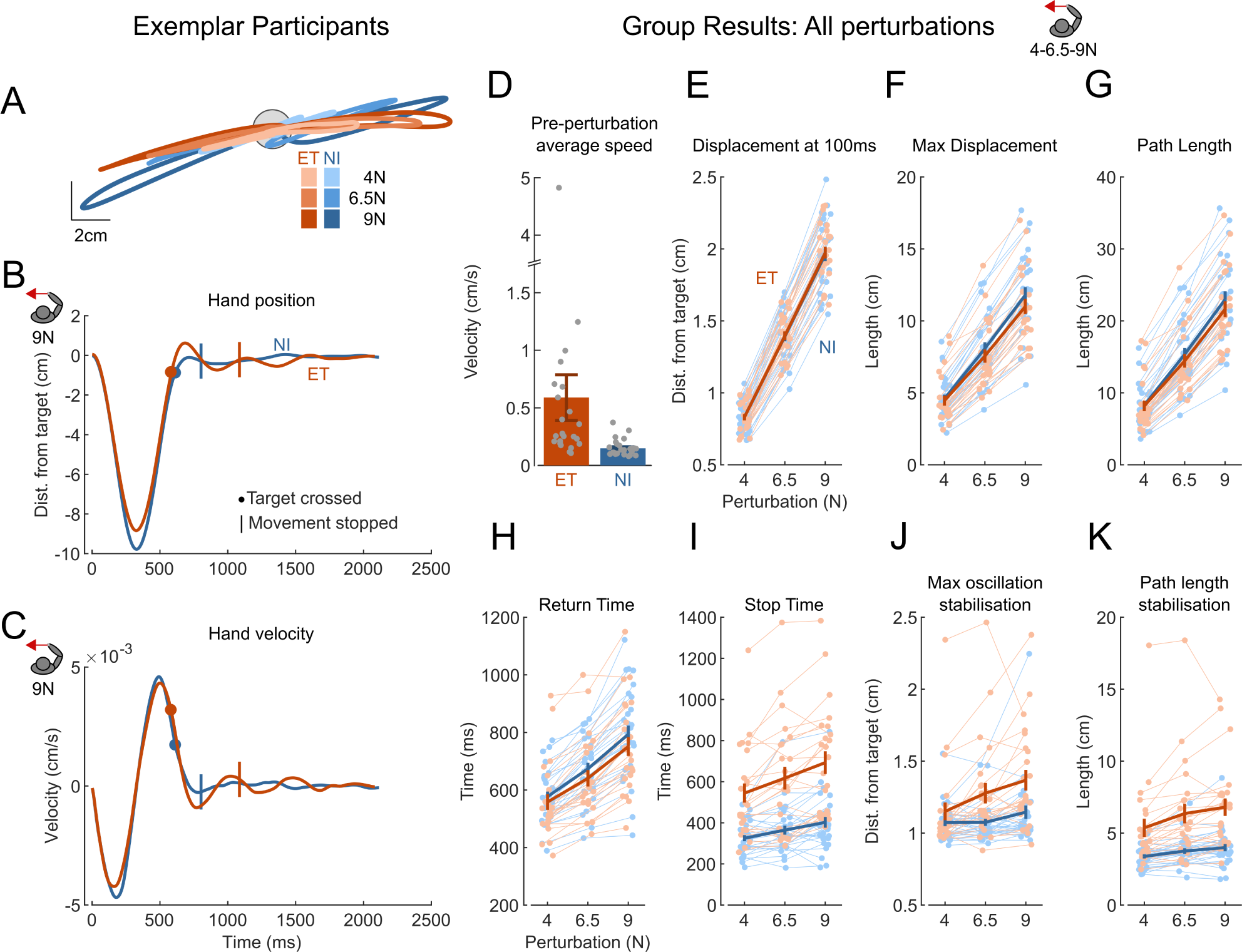
Perturbation *trials, ET patients’ difficulties in the stabilization phase are increased with perturbation magnitude*. **A**, Exemplar participants’ hand path during the last perturbation trial, the 2 analyzed directions and 3 amplitudes are represented. **B**, Distance in the X-axis of the exemplar hand path for the 9 N -x perturbation. **C**, X-Velocity profile of the same trials. **B, C**, Dot: time at which the hand re-entered the target for the first time; Bar: movement stop condition met: velocity&lt;3 cm/s for at least 200 ms (see Methods). **D**, Pre-perturbation average speed, averaged over all perturbation magnitudes, gray dots represent individual behavior. **E-H**, Movement phase, prior first target crossing. **E**, Distance of the hand from the target center at 100 ms. **F**, Maximum displacement, **G**, Path length **H**, Time elapsed between perturbation onset and first target crossing. **I-K**, Stabilization phase. **I**, From target re-entering, time required to stop the hand in the target **J**, Maximum displacement from target center along the X-axis. **K**, Path length. **E-K**, Metrics computed on all -x perturbation trials for each perturbation magnitude. Error bar represents the SEM. Light lines represent individual behavior.

Diagnosis: F(50) =2.18, P=0.03; Diagnosis x Force: F(7743) =4.94, P<10⁻⁴). The interaction between the diagnosis and the force magnitude indicated that performances of the stabilization phase significantly worsened with perturbation amplitude for the three stabilization measures.

In addition to the hand kinematics, we recorded surface muscular activity. We present here recordings of the Pectoralis Major and the Deltoid, respectively the shoulder flexor and extensor stretched by the perturbations aligned with the 𝑥 −axis. No group effects were observed prior to the perturbation (-50-0 ms; LME: Diagnosis: F(50) =-0.89, P=0.38). The initial reflexes elicited by the perturbation were studied and compared between groups shortly after its onset (Figure 3). No significant differences between groups were observed in the short latency window (R1: 25-45 ms; LME: Diagnosis: F(50) =0.11, P=0.91; Force: F(7743) =4.02, P<10^−4^; Diagnosis x Force: F(7743) =1.05, P=0.30) and during R2, the first part of the long-latency window (45-75 ms; LME: Diagnosis: F(50) =0.43, P=0.74; Force: F(7743) =4.02, P<10^−4^; Diagnosis x Force: F(7743) =-0.06, P=0.95). Note that the absence of significant differences between NI and ET in R1 and R2 is not due to an absence of response as we observed a strongly significant effect of the Force on the muscular activity. Differences between the two groups are visible starting from R3, the second part of the long-latency window (75-105 ms; LME: Diagnosis: F(50) =0.73, P=0.47; Force: F(7743) =4.02, P<10^−4^; Diagnosis x Force: F(7743) =-3.66, P<10⁻³).

**Figure 3:**
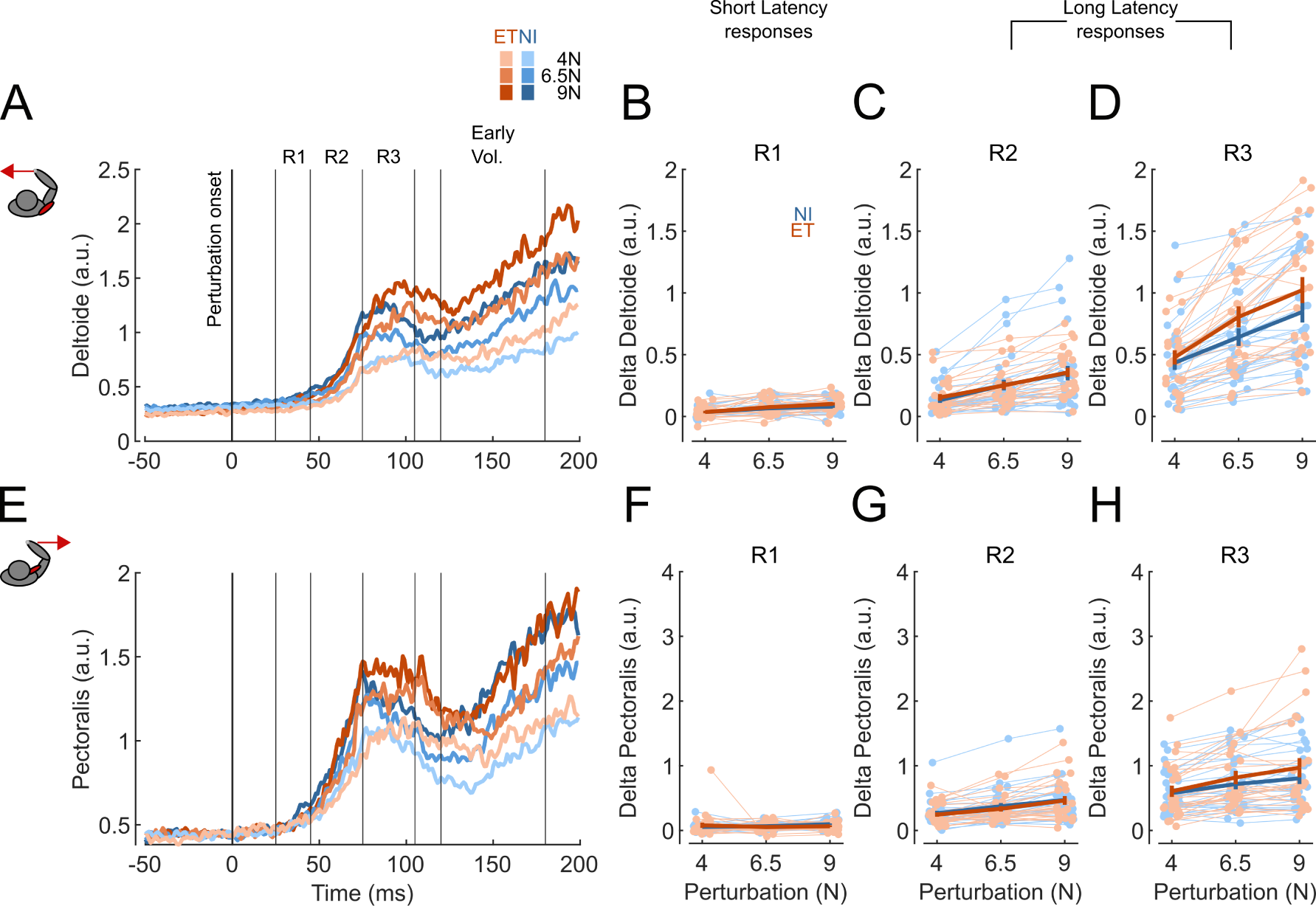
S*h*ort *and Long-Latency responses in the perturbation trials.* **A**, Early deltoid EMG responses averaged over each group for the 3 perturbation magnitudes aligned with negative direction of the 𝑥-axis. Vertical lines delineate the different time windows analyzed. **B-D**, Binned data over the different time windows for the 3 perturbation magnitudes. Average baseline (-50 to 0 ms) was subtracted from the data for illustration purpose. Error bars represent the SEM. Light traces depict individual behavior. **E-H**, Pectoralis response for the perturbations aligned with the positive 𝑥-direction. Same format as A-D.

These results suggest similar initial response following perturbation and highlight intact proprioception in ET. Impairments are visible in the late long-latency responses (LLRs), from 75ms. Surprisingly, this difference was associated with a slightly larger R3 response in the ET group, which was dependent on the perturbation magnitude as indicated by the interaction term. We demonstrate below that this effect is in fact expected in theory, assuming partial delay compensation in ET.

### Computational model of feedback control in ET

In order to describe the behavior of ET patients from a theoretical perspective, we reproduced our experimental observations with a closed loop control model featuring a state estimator that compensates for sensorimotor delays. We modeled a Linear-Quadratic-Gaussian (LQG) controller taking into account delayed sensory feedback (Fig 1C: dotted line) and internal representation of the body dynamics (Fig1C: dashed line) to produce an optimal motor command. The delay in the sensory feedback is compensated by extrapolating the sensory feedback over the delay interval (Fig1C: orange line), which is performed explicitly in this approach to study the impact of errors in this operation.

We considered a scenario in which the sensory feedback delay was underestimated by the controller: the value of the delay used in the controller was 70% of the system’s delay (𝛿𝑡 = 55 𝑚𝑠, ^𝛿^^𝑡 = 0.7𝛿𝑡). This model previously showed that a mismatch in the state estimator associated with long-latency delays could also produce oscillations (Crevecoeur & Gervers, 2019).

In such case, the simulations for which the delay was underestimated produced trajectories bearing striking similarities with those of ET participants. Initially, the arm angle and angular velocity matched the simulations with no errors in delay estimation. We can notice a slightly smaller maximum hand deviation in the underestimated case, a trend that was also observed in the experimental results for the ET population (Figure 2F). An overshoot was observed when bringing the hand back to the target and was followed by oscillations during the stabilization phase (Figure 4B and C). Oscillations amplitude scaled with perturbation magnitude and their frequency matched the oscillations observed with our participants; with a higher power spectral density for ET patients in the range 0 to 15Hz (Figure 4D), and a peak frequency located around 4Hz as in the data. The motor commands produced by the model during the LLR window showed no difference during R2, and early differences in R3, similarly to what was observed between NI and ET participants (Figure 4E). This effect was robust as the presence of oscillations and increased gains persisted even when varying parameters such as delay and inertia. Simulations with parameters leading to the most extreme changes of slopes are visible on Figure 4E.

**Figure 4:**
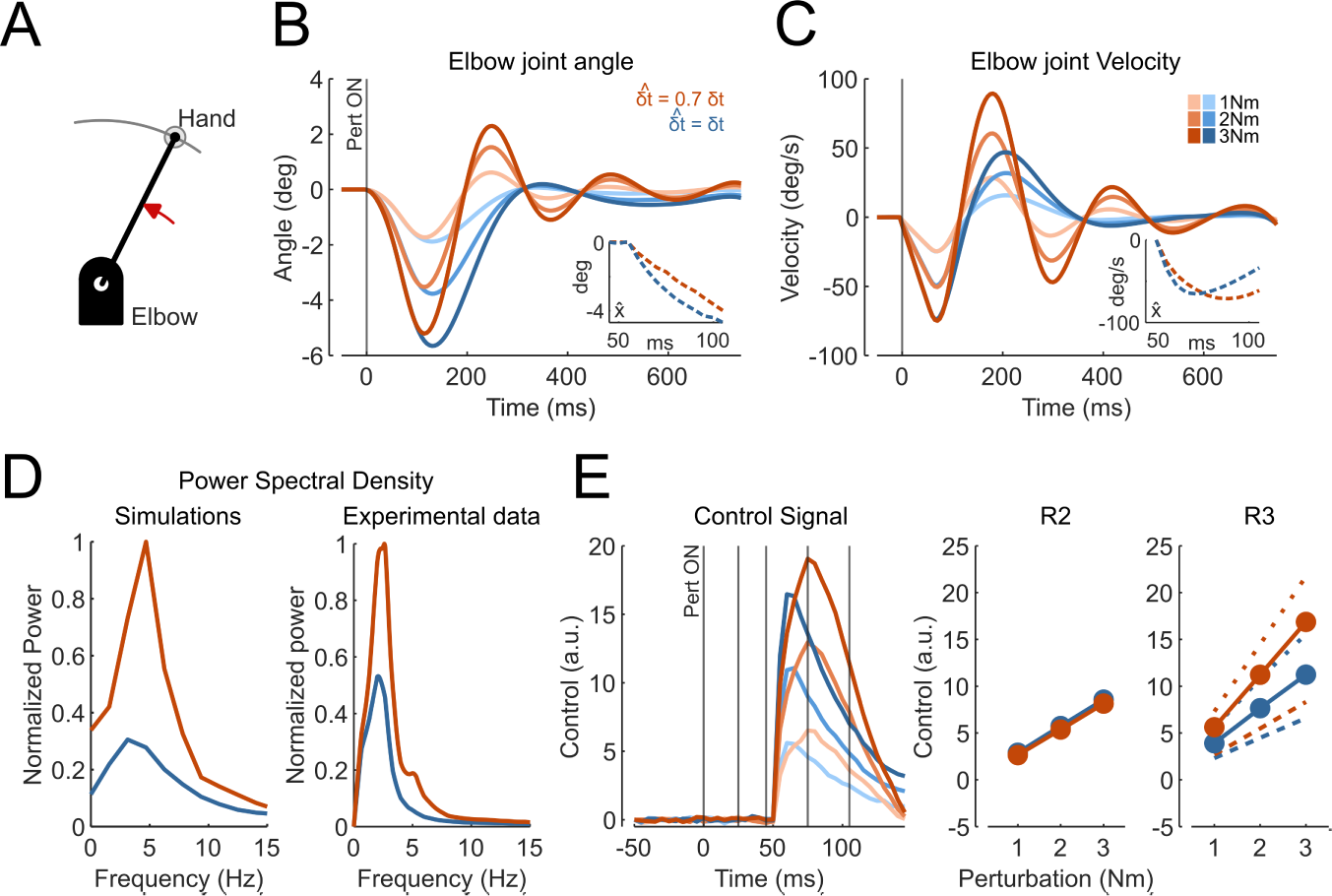
C*o*mputational *model of feedback control in ET.* A, Simulations of step perturbations in a single joint system. B-E, Model underestimating the sensory delays (^𝛿^^𝑡 = 0.7𝛿𝑡, orange) and model with no delay estimation error (𝛿𝑡 = 55 𝑚𝑠 , blue). B, Elbow joint angle following perturbation (1, 2 or 3 Nm). Insets represent the estimated state over the long-latency windows for the larger perturbation. C, Elbow joint angular velocity following perturbation. Inset same as B. D, Average normalized power spectral density across all perturbations for the simulated (left) and experimental (right) data. E, (Left) Average control signal for the 3 perturbation magnitudes ([1, 2, 3] Nm). Detail of the R2 (45-75 ms, middle panel) and R3 (75-105 ms, right panel) long-latency Windows. Additional traces on the right panel represent the most extreme slope changes following changes in delay and inertia values. Lower limit: *I*= 0.2 Kgm², 𝛿𝑡 = 45 ms. Higher limit: *I*=0.1 Kgm²,𝛿𝑡 = 65ms.

We then assessed the sensitivity of the model properties as a function of different parameters. First, we observed that the main frequency of the oscillations remained unaffected when varying the limb inertia (Figure 5A), which reminds ET as tremor frequency is also unaffected by arm loading (Elble et al., 1994). Similarly, varying the delay or delay error had an effect on the amplitude of the oscillations, while keeping the same tremor frequency (Figure 5 B&C). To identify the origin of these oscillations, we separated the contribution of a delay estimation error solely in the extrapolation of the state (first term of equation 11) with an error solely in the extrapolation of motor commands (second term of eq. 11). We observed that the oscillations mainly arose from errors of the latter component (Figure 5D-G). These results confirm the possibility that the initial pass through the state estimator produced comparable responses, as reflected by intact R2 responses, but errors accumulated over time due to the underestimation of the delay in the extrapolation of motor commands, resulting in oscillations during the later phases of the corrective movement.

**Figure 5:**
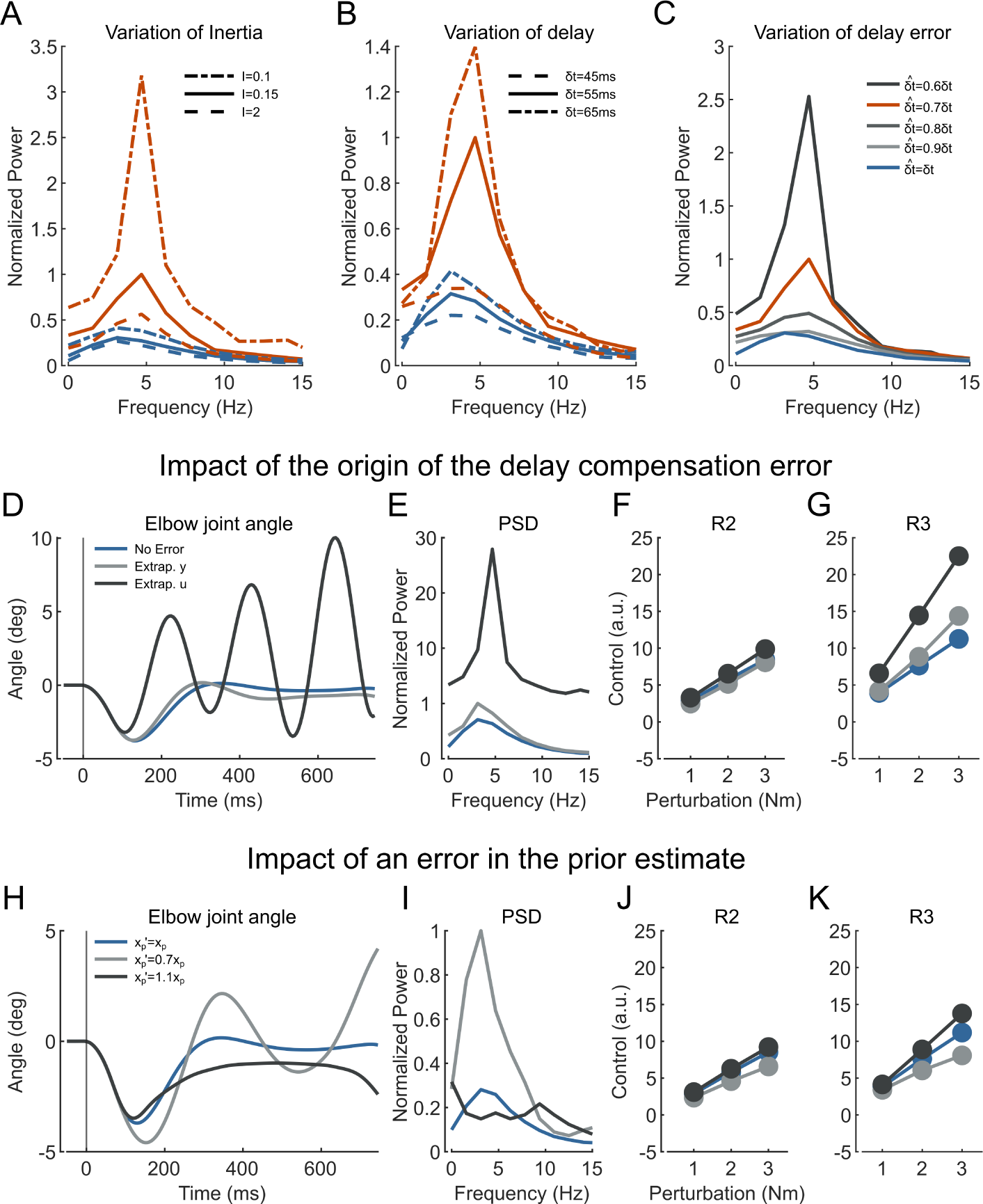
Sensitivity analyses of our feedback control model of ET. A, Variation of PSD under different values of inertia for the model underestimating the sensory delays (^𝛿^^𝑡 = 0.7𝛿𝑡, orange) and the model with no delay estimation error (𝛿𝑡 = 55 𝑚𝑠 , blue). B, same as A for different values of delay (𝛿𝑡 = 45𝑚𝑠/55𝑚𝑠/65𝑚𝑠). C, Effect of different delay errors on the PSD. D-G, Impact of the origin of the delay estimation error. Three cases were considered: an error (^𝛿^^𝑡 = 0.7𝛿𝑡) solely in the extrapolation of the sensory feedback over the delay interval (light grey), an error (^𝛿^^𝑡 = 0.7𝛿𝑡) solely in the extrapolation of the motor commands (dark grey), or no errors (^𝛿^^𝑡 = 𝛿𝑡) in delay estimation (blue). D, Elbow joint angle following a 2Nm perturbation. E, Average normalized power spectral density across all perturbations. F-G, Average control signal for 3 perturbation magnitudes ([1, 2, 3] Nm). F, Detail of the R2 (45-75 ms) and G, R3 (75-105 ms) Long Latency Windows. H-I, Impact of an error in the prior estimate 𝑥_𝑝_. Three cases considered: underestimation of the prior estimate (𝑥^′^ = 0.7𝑥_𝑝_, light gray), overestimation of the prior estimate (𝑥_𝑝_′ = 1.1𝑥_𝑝_, dark grey) or no error (𝑥_𝑝_′ = 𝑥_𝑝_, blue). Subpanels same as D-G.

All simulations above considered a static error in the delay parameter. To investigate the effect of a noisy error in the delay, we considered a delay composed of a fixed, or mean error, plus a random error drawn from uniform distribution (see Methods). Interestingly, simulations exhibited similar oscillations around the target, and an increase in the early response component compatible with the increase measured in R3. In contrast, a random delay error in the absence of a mean error did not produce oscillations. Thus, it seemed that the mean of the error was more important than a random source of the error, however a fuller treatment of random delay errors with more general sources of noise is needed in future studies.

Finally, we investigated the possibility that oscillations arose from errors in other parameters of the model and discuss their link to the different pathways associated with state estimation. First, an up- or down-scaling of the Kalman gains accelerated or slowed down the model response to the perturbation, but it did not produce any oscillation. Then, we introduced an error in the prior estimate, 𝑥_𝑝_ (Eqn. 10, Figure 5 H-K). When an overestimation was made, it reproduces a slight increase in the R3 response, as observed in experimental data. However, it did not lead to oscillations and while the system was unable to reach back to the target (Fig. 5H). We noted that testing higher values of multiplicative errors produced numerical instability (the scaling factor was 1.1). Conversely, underestimating the prior produced oscillations at a lower frequency than observed in ET, and was associated with a decrease in the initial response, which was incompatible with our experimental observations. Finally, we introduced errors in the internal models underestimating 𝐴_𝑑_ or 𝐵_𝑑_ following previous work (Takei et al., 2021). An error in these parameters produced effects that were generally not compatible with the data, except for 𝐵_𝑑_ which impacted the same summation as the delay (Eqn. 11). As these parameters impact all operations, we privileged the delay error that had a more specific role in the feedback pathway.

To summarize, based on oscillations and the transient increase in control responses observed in R3 for patients, we conclude that an erroneous delay estimation appears was the only hypothesis that reproduced all experimental features observed in ET patients while varying a single parameter.

## Discussion

We investigated the hypothesis that ET originates from an erroneous compensation of the sensory delays. With a postural perturbation task, we observed feedback control impairments in the ET population. The initial impact of the perturbation was similar for the ET and NI volunteers, in the kinematics as well as the initial stretch responses. However, ET patients experienced perturbation dependent deficits when stabilizing their hand in the target, and they exhibited an increased R3 long-latency response suggesting errors in the feedback controller. We reproduced these results with a computational model erroneously compensating for long-latency delays. The simulations were performed with a value of the delay that was smaller than the true delay, producing an intact initial response to the perturbation followed by an accumulation of errors over time that led to impaired feedback control and to the emergence of oscillations that scaled with the force magnitude and matched the frequency of oscillations measured in ET patients.

To identify which pathway is affected in ET, we can divide our computational model into 3 main processes: 1- the processing of the afferent sensory information, 2- the prior estimate of the state of the system and of the environment (such as the external torque, estimated), 3- and finally the delay compensation pathway requiring internal monitoring of the effect of motor commands during the delay interval. Our results suggest that the sensory feedback is intact in ET. Indeed, because we did not observe any difference in short-latency response, and early long-latency responses (R2), we can conclude that the peripheral system and the early components of the stretch responses are preserved in ET. This result is also consistent with a recent study reporting intact proprioception in ET patients (Butler et al., 2023). Regarding the prior estimate of the state, simulations revealed that introducing an error that over or underestimate the prior did not reproduce the oscillations coupled with the increased R3 response observed in ET (Fig 5 H-K). Additionally, in our previous study, we reported motor adaptation impairments in ET. The anticipation of the perturbation was preserved while the metrics related to motor control were impaired (Blondiaux et al., 2023). Their ability to anticipate an external disturbance suggests the preservation of patients’ ability to form an internal prediction of their movement even in the presence of a perturbation.

Thus, on the one hand, it is possible that both internal priors and sensory feedback are preserved, while, on the other hand, we observed clear deficits that could be linked to the feedback controllers as they scaled with the perturbation magnitude. The third model component, linked to the internal monitoring of motor commands, becomes a unique candidate pathway to account for sensorimotor deficits in this population: it is clearly a component of the state estimator, and therefore, of the feedback controller, and can be dissociated from both the sensory system and the formation of internal priors. Moreover, a selective alteration of the operation performed in this component produced all features observed experimentally, namely preserved R2 response, increased R3 response and impaired stabilization scaling with perturbation magnitude. However, it is important to note that, while an error in delay estimation is also affecting the pathway processing the sensory feedback, we observed that errors solely in this pathway does not lead to the emergence of oscillations. Consequently, we concluded that the impairment was attributable to the erroneous integration of motor commands. A clear added value of our modeling work is to provide a quantitative model of ET based on the simple assumption that their feedback control is not well adjusted to the true value of long-latency delays.

Considering some quantitative differences between our results and the simulations, a note of caution is warranted. First, an approximation of body dynamics was used in our model for the derivation of the controller. The simulations reported in this study are a simplified version of the task as we only modeled single-joint movements around a fixed axis. Our participants performed movements in the 3D space using the handle of the robotic device and therefore responded to the perturbation using more than one joint. In addition, the muscular activity collected during the experiment and the control signal extracted from the simulations did not fully match in terms of temporal profiles. Indeed, the model did not take the details of non-linear muscle dynamics into account nor the effect of antagonist muscles. In spite of these differences, the range of oscillations frequencies was similar across simulations and data, suggesting that the net effect of the muscle responses may correspond to the simulated control signal.

Despite these simplifications, we did observe an increase of the R3 responses both in the experimental data and the simulations, an effect that remained present even when varying several simulation parameters such as the limb inertia, the value of the delay or delay error. While the estimation of the perturbation magnitude was intact, simulations revealed that the velocity was overestimated starting already ∼70 ms following the load onset (Figure 4 B & C) which resulted in an increased gain in the response despite a small underestimation of the position.

Based on these results, a next challenge is to identify the neural pathways behind these operations to better identify the origin of the deficits in ET. To date cerebellum is one of the main candidates. Traditional views of the role of cerebellum in motor control have considered that this region was involved in forward models (Kawato, 1999). However, it has been also associated with state estimation (Diedrichsen & Bastian, 2014; Franklin & Wolpert, 2011; Miall et al., 1993; Wolpert et al., 1998), which is a component of feedback control models and of LLR (Crevecoeur & Scott, 2013; Todorov & Jordan, 2002). We also know from human and animal studies that a suboptimal state estimation leads to incorrect movements or oscillatory behavior (Flament et al., 1984; Miall et al., 2007; Takei et al., 2021). Interestingly, limb-imposed displacement induces changes in cerebellar nuclei activity within 70 ms, aligning with the timing of the long-latency pathway (Strick, 1983). In addition, severe cerebellar dysfunction induced by cerebellar ataxia or dentate nucleus cooling in monkeys is known to reduce drastically LLRs responses (Kurtzer et al., 2013; Meyer-Lohmann et al., 1975), making this structure a contributor to long-latency feedback pathway. Thus, the functional implication of cerebellum in LLR, and the possibility that the corresponding operation corresponds to state estimation, is in our view the candidate mechanism that is potentially altered in ET, while other cerebellar functions linked to forward models and anticipation may be preserved in this population (Blondiaux et al., 2023).

We suggest that instead of a severe cerebellar disruption leading to a substantial response attenuation as reported in cerebellar ataxia, ET could originate from subtle and partial cerebellar dysfunction such as a small mismatch in delay estimation that is enough to produce sensorimotor deficits. This error in delay estimation does not affect the initial pass of the sensory feedback through the state estimator as observed with the intact initial responses, compatible with the observation that peripheral and spinal contributions seemed intact (R1 and R2). Yet, the accumulation of errors with multiple passes through the state estimator generated oscillations. Future studies should be performed to relate this functional hypothesis with the anatomical structures behind it using for example neuroimaging. Interestingly, the link between state estimation and cerebellum as well as its involvement in LLRs aligns with numerous indicators of a cerebellar origin in ET as highlighted by neuroimaging studies (Holtbernd & Shah, 2021; Mavroudis et al., 2019; Pietracupa et al., 2021; Tikoo et al., 2020), as well as postmortem studies (Louis et al., 2007; Louis & Faust, 2020a) which report pathological cerebellar changes in ET.

Overall, this study reported intact short and initial long-latency responses in ET during a postural perturbation task. However, perturbation-dependent deficits were observed when ET participants stabilized their hand in the home target, confirming impairments in feedback control. Based on computational modeling, we suggest that these deficits may stem from an erroneous compensation for sensorimotor delays, attributed to an underestimation of the delay by the controller.

## Acknowledgments

Florence Blondiaux is a FRIA grantee of the Fonds de la Recherche Scientifique – FNRS (Be). Lise Colmant is a Research Fellow of the Fonds de la Recherche Scientifique – FNRS (Be). The FNRS provided salary and research support for Bernard Hanseeuw under grants n° CCL40010417 and n° FRFS-WELBIO40010035. Frédéric Crevecoeur is supported by a grant from the FNRS under grant number 1.C.033.18.

## References

1. Bhatia, K. P., Bain, P., Bajaj, N., Elble, R. J., Hallett, M., Louis, E. D., Raethjen, J., Stamelou, M., Testa, C. M., Deuschl, G., & the Tremor Task Force of the International Parkinson and Movement Disorder Society. (2018). Consensus Statement on the classification of tremors. From the task force on tremor of the International Parkinson and Movement Disorder Society. Movement Disorders, *33*(1), 75–87. 10.1002/mds.27121

2. Blondiaux, F., Lebrun, L., Hanseeuw, B. J., & Crevecoeur, F. (2023). Impairments of saccadic and reaching adaptation in Essential Tremor are linked to movement execution. Journal of Neurophysiology, jn.00165.2023. 10.1152/jn.00165.2023

3. Butler, A. A., Diong, J., Lidman, K., Adler, J., Wardman, D. L., Gandevia, S. C., & Héroux, M. E. (2023). Upper Limb Function but Not Proprioception is Impaired in Essential Tremor: A Between-Groups Study and Causal Mediation Analysis. Tremor and Other Hyperkinetic Movements, 13(1), 1. 10.5334/tohm.731

4. Crevecoeur, F., & Gervers, M. (2019). Filtering compensation for delays and prediction errors during sensorimotor control. 2954, 2925–2954. 10.1162/NECO

5. Crevecoeur, F., Kurtzer, I., & Scott, S. H. (2012). Fast corrective responses are evoked by perturbations approaching the natural variability of posture and movement tasks. Journal of Neurophysiology, 107(10), 2821–2832. 10.1152/jn.00849.2011

6. Crevecoeur, F., & Scott, S. H. (2013). Priors Engaged in Long-Latency Responses to Mechanical Perturbations Suggest a Rapid Update in State Estimation. PLoS Computational Biology, 9(8). 10.1371/journal.pcbi.1003177

7. Crevecoeur, F., & Scott, S. H. (2014). Beyond Muscles Stiffness: Importance of State-Estimation to Account for Very Fast Motor Corrections. PLoS Computational Biology, 10(10), e1003869. 10.1371/journal.pcbi.1003869

8. De Comite, A., Crevecoeur, F., & Lefèvre, P. (2021). Online modification of goal-directed control in human reaching movements. Journal of Neurophysiology, 125(5), 1883–1898. 10.1152/jn.00536.2020

9. Diedrichsen, J., & Bastian, A. J. (2014). Cerebellar Function. The Cognitive Neurosciences, 451.

10. Elble, R. J. (2003). Characteristics of physiologic tremor in young and elderly adults. Clinical Neurophysiology, 114(4), 624–635. 10.1016/S1388-2457(03)00006-3

11. Elble, R. J., Higgins, C., Leffler, K., & Hughes, L. (1994). Factors influencing the amplitude and frequency of essential tremor. Movement Disorders, 9(6), 589–596. 10.1002/mds.870090602

12. Fahn, S., Tolosa, E., & Marin, C. (1993). Clinical Rating Scale for Tremor. Parkinson’s Disease and

13. *Movement Disorders*, *2*, 271–280.

14. Flament, D., Vilis, T., & Hore, J. (1984). Dependence of cerebellar tremor on proprioceptive but not visual feedback. Experimental Neurology, 84(2), 314–325. 10.1016/0014-4886(84)90228-0

15. Franklin, D. W., & Wolpert, D. M. (2011). Computational Mechanisms of Sensorimotor Control. Neuron, 72(3), 425–442. 10.1016/j.neuron.2011.10.006

16. Fridman, E. (2014). Introduction to Time-Delay Systems: Analysis and Control. Springer International Publishing. 10.1007/978-3-319-09393-2

17. Holtbernd, F., & Shah, N. J. (2021). Imaging the Pathophysiology of Essential Tremor—A Systematic Review. Frontiers in Neurology, 12, 680254. 10.3389/fneur.2021.680254

18. Ibrahim, M. F., Beevis, J. C., & Empson, R. M. (2021). Essential Tremor – A Cerebellar Driven Disorder? Neuroscience, 462(November), 262–273. 10.1016/j.neuroscience.2020.11.002

19. Kawato, M. (1999). Internal models for motor control and trajectory planning. Current Opinion in Neurobiology, 9(6), 718–727. 10.1016/S0959-4388(99)00028-8

20. Kurtzer, I., Trautman, P., Rasquinha, R. J., Bhanpuri, N. H., Scott, S. H., & Bastian, A. J. (2013). Cerebellar damage diminishes long-latency responses to multijoint perturbations. Journal of Neurophysiology, 109(8), 2228–2241. 10.1152/jn.00145.2012

21. Louis, E. D., & Faust, P. L. (2020a). Essential tremor pathology: Neurodegeneration and reorganization of neuronal connections. Nature Reviews Neurology, 16(2), 69–83. 10.1038/s41582-019-0302-1

22. Louis, E. D., & Faust, P. L. (2020b). Essential Tremor Within the Broader Context of Other Forms of Cerebellar Degeneration. The Cerebellum, 19(6), 879–896. 10.1007/s12311-020-01160-4

23. Louis, E. D., Faust, P. L., Vonsattel, J.-P. G., Honig, L. S., Rajput, A., Robinson, C. A., Rajput, A., Pahwa, R., Lyons, K. E., Ross, G. W., Borden, S., Moskowitz, C. B., Lawton, A., & Hernandez, N. (2007). Neuropathological changes in essential tremor: 33 cases compared with 21 controls. Brain, 130(12), 3297–3307. 10.1093/brain/awm266

24. Louis, E. D., & Ferreira, J. J. (2010). How common is the most common adult movement disorder? Update on the worldwide prevalence of essential tremor. Movement Disorders, 25(5), 534–541. 10.1002/mds.22838

25. Mavroudis, I., Petridis, F., & Kazis, D. (2019). Neuroimaging and neuropathological findings in essential tremor. Acta Neurologica Scandinavica, 139(6), 491–496. 10.1111/ane.13101

26. Meyer-Lohmann, J., Conrad, B., Matsunami, K., & Brooks, V. B. (1975). Effects of dentate cooling on precentral unit activity following torque pulse injections into elbow movements. Brain Research, 94(2), 237–251. 10.1016/0006-8993(75)90059-1

27. Miall, R. C., Christensen, L. O. D., Cain, O., & Stanley, J. (2007). Disruption of state estimation in the human lateral cerebellum. PLoS Biology, 5(11), 2733–2744. 10.1371/journal.pbio.0050316

28. Miall, R. C., Weir, D. J., Wolpert, D. M., & Stein, J. F. (1993). Is the Cerebellum a Smith Predictor? Journal of Motor Behavior, 25(3), 203–216. 10.1080/00222895.1993.9942050

29. Mondie, S., & Michiels, W. (2003). Finite spectrum assignment of unstable time-delay systems with a safe implementation. IEEE Transactions on Automatic Control, 48(12), 2207–2212. 10.1109/TAC.2003.820147

30. Pietracupa, S., Bologna, M., Tommasin, S., Berardelli, A., & Pantano, P. (2021). The Contribution of Neuroimaging to the Understanding of Essential Tremor Pathophysiology: A Systematic Review. Cerebellum, 0123456789. 10.1007/s12311-021-01335-7

31. Pruszynski, J. A., & Scott, S. H. (2012). Optimal feedback control and the long-latency stretch response. Experimental Brain Research, 218(3), 341–359. 10.1007/s00221-012-3041-8

32. Scott, S. H. (2016). A Functional Taxonomy of Bottom-Up Sensory Feedback Processing for Motor Actions. Trends in Neurosciences, 39(8), 512–526. 10.1016/j.tins.2016.06.001

33. Shanker, V. (2019). Essential tremor: Diagnosis and management. The BMJ, 366. 10.1136/bmj.l4485

34. Stein, R. B., & Oĝuztöreli, M. N. (1976). Tremor and other oscillations in neuromuscular systems. Biological Cybernetics, 22(3), 147–157. 10.1007/BF00365525

35. Strick, L. (1983). The influence of motor preparation on the response of cerebellar neurons to limb displacements. The Journal of Neuroscience, 3(10).

36. Takei, T., Lomber, S. G., Cook, D. J., & Scott, S. H. (2021). Transient deactivation of dorsal premotor cortex or parietal area 5 impairs feedback control of the limb in macaques. Current Biology, 31(7), 1476–1487.e5. 10.1016/j.cub.2021.01.049

37. Tikoo, S., Pietracupa, S., Tommasin, S., Bologna, M., Petsas, N., Bharti, K., Berardelli, A., & Pantano, P. (2020). Functional disconnection of the dentate nucleus in essential tremor. Journal of Neurology, 0123456789. 10.1007/s00415-020-09711-9

38. Todorov, E., & Jordan, M. I. (2002). Optimal feedback control as a theory of motor coordination. Nature Neuroscience, 5(11), 1226–1235. 10.1038/nn963

39. Trillenberg, P., Führer, J., Sprenger, A., Hagenow, A., Kömpf, D., Wenzelburger, R., Deuschl, G., Heide, W., & Helmchen, C. (2006). Eye-hand coordination in essential tremor: Eye-Hand Coordination in Essential Tremor. Movement Disorders, 21(3), 373–379. 10.1002/mds.20729

40. Wolpert, D. M., Miall, R. C., & Kawato, M. (1998). Internal models in the cerebellum. Trends in Cognitive Sciences, 2(9), 338–347. 10.1016/S1364-6613(98)01221-2

